# New single nucleotide polymorphisms (SNPs) in homologous recombination repair genes detected by microarray analysis in Polish breast cancer patients

**DOI:** 10.1101/088948

**Authors:** Hanna Romanowicz, Dominik Strapagiel, Marcin Słomka, Marta Sobalska-Kwapis, Ewa Kępka, Anna Siewierska-Górska, Marek Zadrożny, Beata Smolarz

## Abstract

**Purpose of the study:** Breast cancer is the most common cause of malignancy mortality in women worldwide. This study aimed at localising homologous recombination repair (HR) genes and their chromosomal loci and correlating their nucleotide variants with susceptibility to breast cancer. In this study authors analysed the association between single nucleotide polymorphisms (SNPs) in homologous recombination repair genes and the incidence of breast cancer in the population of Polish women.

**Methods:** Blood samples from 94 breast cancer patients were analysed as test group. Individuals were recruited into the study at the Department of Oncological Surgery and Breast Diseases of the Institute of the Polish Mother’s Memorial Hospital in Lodz, Poland. Healthy controls (n=500) were obtained from the Biobank Laboratory, Department of Molecular Biophysics, University of Lodz. Then, DNA of breast cancer patients was compared with one of disease-free women. The test was supported by microarray analysis.

**Results:** Statistically significant correlations were identified between breast cancer and 3 not described previously single nucleotide polymorphisms (SNPs) of homologous recombination repair genes *BRCA1* and *BRCA2*: rs59004709, rs4986852 and rs1799950.

**Conclusions:** Further studies on larger groups are warranted to support the hypothesis of correlation between the above-mentioned genetic variants and breast cancer risk.

## Introduction

Breast cancer is one of the most common malignancies in females and its morbidity is still growing [1, 2]. This phenomenon is particularly visible in developed societies what can be scientifically explained by the rising prevalence of common risk factors of breast cancer such as: obesity, hormone replacement therapy and alcohol consumption in these populations. Moreover, the positive correlation between the abovementioned conditions and the risk of breast carcinoma is one of the mostly proven clinical interactions in entire oncology.

Besides clinical features that predispose towards developing breast cancer, the study of DNA has also given clear evidence of links between genetic variations and this malignancy.

The genes of DNA lesion repair system play the key role in maintaining genome integrity and in controlling the repair of mutation-affected DNA [3-8]. Malfunction of these genes would result in rapid accumulation of errors within DNA, which would eventually make further cell survival impossible. Moreover, proper DNA repair system ensures genomic integrity and plays a significant role in its protection against effects of carcinogenic factors [3-8]. The variability of DNA repair genes may also carry clinical significance when correlated with the risk of development of specific types of cancer, rules of their prophylactics and possible therapy [7, 8].

The DNA repair process usually encompasses two stages: excision of affected DNA lesion and then its repair synthesis. This is how the system performs through base-excision repair (BER), nucleotide excision repair (NER) and mismatch repair (MMR). Totally converse is the repair system activity by direct lesion reversal, in which there is merely a single-stage process with maintained integrity of the DNA phosphodiester chain and the system of recombination repair (HR).

Literature data suggests that genomic rearrangements are frequent in breast cancer cells [9-12]. Moreover, these phenomena are believed to result from an aberrant repair of DNA double-strand breaks (DSBs). These even may bring a loss of some chromosomes, causing translocation of genetic material between them. If not repaired, these genetic disorders may lead to down-regulation of transcription and further on to development of various malignancies [13, 14]. The repair by recombination itself enables the removal of numerous serious DNA lesions, including even the double-stranded breaks [15, 16].

In Caucasian population the variability of DSBs repair genes may play a role in breast cancer risk [17]. Polymorphisms in DNA repair genes may alter the activity of acting proteins and determine cancer susceptibility in this very fashion [18]. State-of-the-art research focuses on the analysis of versatile genetic aspects and on the attempt to associate ones with clinical manifestation of carcinogenesis. Large effort has been lately put into investigation of single nucleotide polymorphisms (SNPs), which may underline the differences in ones susceptibility and natural history of diseases.

In this study we analysed the association between breast cancer incidence and the SNPs encountered in six HR-related genes: *XRCC2*, *XRCC3, RAD51, RAD52, BRCA1* and *BRCA2*.

## Materials and methods

### Patients

The test group comprised 94 blood samples obtained from breast cancer patients treated in the Department of Oncological Surgery and Breast Diseases, Polish Mother’s Memorial Hospital in Lodz. 500 DNA samples collected from unrelated disease free women selected randomly from Polish female population of the database of the Biobank Lab, Department of Molecular Biophysics, University of Lodz, served as controls [19]. Both patients and controls were Caucasians of equivalent ethnic and geographical origins. The demographic data and pathological features of the patients are summarized in Table 1. All breast cancer samples were staged accordingly to Scarf-Bloom-Richardson criteria. Blood samples were derived in EDTA-vacuum tubes, and preserved frozen in −20 °C until initial processing commenced. DNA was automatically isolated from 200 μl of blood using MagNA Pure LC 2.0 Instrument (Roche Applied Science, Indianapolis, IN, USA) and MagNA Pure LC DNA Isolation Kit I, according to manufacturer’s instructions. The High Performance II protocol was launched during the isolation protocol. Elution volume was 100 μl. DNA was quantified using broad range Quant-iT^TM^ dsDNA Broad Range Assay Kit (Invitrogen^TM^, Carlsbad, CA, USA). All DNA samples were subject of quality control in PCR reaction for sex determination [20]. Afterwards, all DNA samples included in the study were diluted to 50 ng/μl in sterile DNase free water, the concentration was then reconfirmed using the same fluorometric method. All study participants gave an informed written consent. A formal consent was also issued by the Bioethical Committee of the Institute of the Polish Mother’s Memorial Hospital in Lodz (Approval number, 10/2012) and Review Board at the University of Lodz (Approval number 7/KBBN-UŁ/II/2014).

**Table 1.**
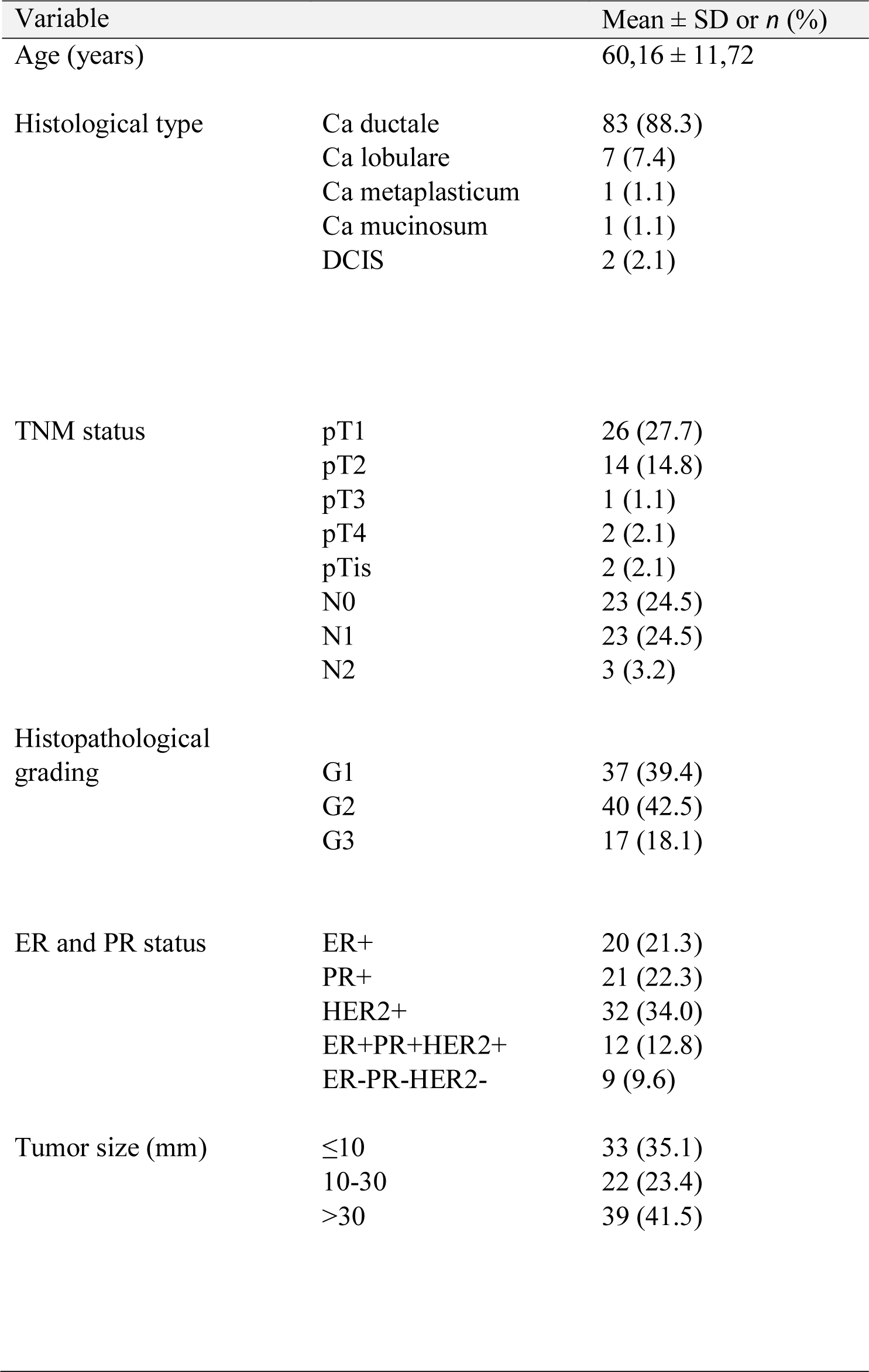
The clinicopathological characteristics of 94 patients with breast cancer.

| Variable | | Mean ± SD or *n* (%) |
| --- | --- | --- |
| Age (years) | | 60,16 ± 11,72 |
| Histological type | Ca ductale | 83 (88.3) |
|  | Ca lobulare | 7 (7.4) |
|  | Ca metaplasticum | 1 (1.1) |
|  | Ca mucinosum | 1 (1.1) |
|  | DCIS | 2 (2.1) |
| TNM status | pT1 | 26 (27.7) |
|  | pT2 | 14 (14.8) |
|  | pT3 | 1 (1.1) |
|  | pT4 | 2 (2.1) |
|  | pTis | 2 (2.1) |
|  | N0 | 23 (24.5) |
|  | N1 | 23 (24.5) |
|  | N2 | 3 (3.2) |
| Histopathological grading | G1 | 37 (39.4) |
|  | G2 | 40 (42.5) |
|  | G3 | 17 (18.1) |
| ER and PR status | ER+ | 20 (21.3) |
|  | PR+ | 21 (22.3) |
|  | HER2+ | 32 (34.0) |
|  | ER+PR+HER2+ | 12 (12.8) |
|  | ER-PR-HER2- | 9 (9.6) |
| Tumor size (mm) | $\leq 10$ | 33 (35.1) |
|  | 10-30 | 22 (23.4) |
|  | >30 | 39 (41.5) |

### Genotyping

A total of 94 (test group) and 500 (controls) DNA samples were genotyped for 551495 single nucleotide polymorphisms (SNPs) using an Illumina 24x1 Infinium HTS Human Core Exome (Illumina, Inc., San Diego, CA, USA) according to protocols provided by the manufacturer. DNA samples were amplified which was then followed by enzymatic fragmentation and hybridization to BeadChips. Afterwards, the BeadChips underwent extension and X-staining processes. Then BeadChips were scanned using an iScan (Illumina, Inc., San Diego, CA, USA).

### Sample Quality Control and Statistics

Raw fluorescence intensities were imported to the Genome Studio (v2011.1) with the Genotyping Module. All the data was firstly subject of stringent quality control, including sample exclusion if the call rate was below 0.94 and 10 % and GenCall parameter was below 0.4. We filtered the data from the chromosomal region of six HR-related genes: *XRCC2*, *XRCC3, RAD51, RAD52, BRCA1* and *BRCA2* (Table 2), which gave us 464 SNPs for the analysis. All sequence coordinates in this study are followed by GRCh37/hg19 reference sequence and were obtained from GenBank (http://www.ncbi.nlm.nih.gov). Those SNPs were further tested and they passed visual inspection for cluster quality and their correctness was eventually confirmed. Then, results were exported from Genome Studio using PLINK Input Report Plug-in v2.1.3 by a forward strand. The statistical analysis was performed using the PLINK v2.050 software (http://pngu.mgh.harvard.edu/purcell/plink/) [21].

**Table 2.**
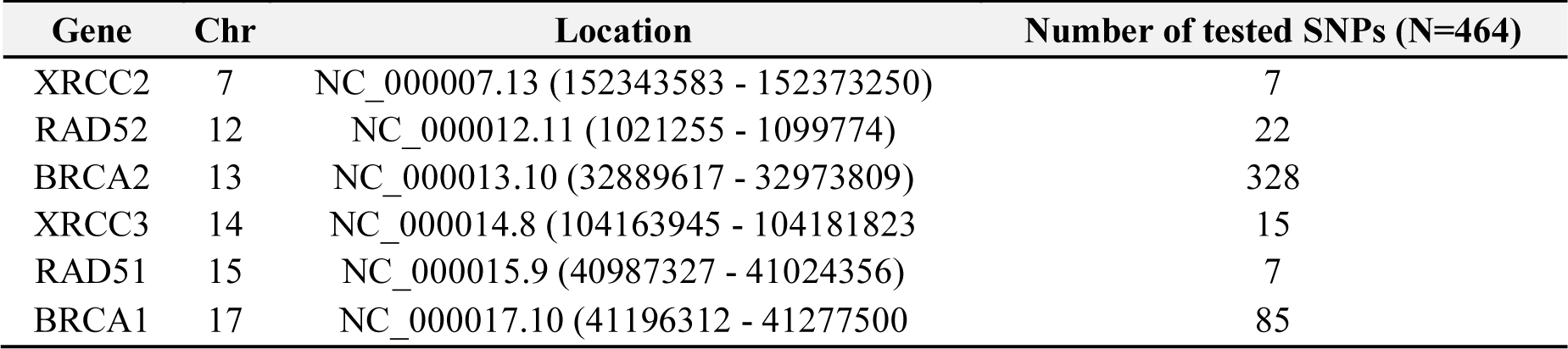
Chromosomal localization of tested HR-related genes and the number of SNPs tested in this study.

All of the tested SNPs (N=464) were first analyzed for consistency with the Hardy– Weinberg exact test. One SNP, rs4987208 *(RAD52),* showed evidence for deviations from Hardy-Weinberg equilibrium (HWE; p< 10^−4^) and was excluded from analysis. According to the estimated minor allele frequencies (MAFs) we excluded from association analysis those SNPs with MAF −0.01 (n=425). Than full genotypic case-control association analysis (CI = 0,95) was performed for 38 SNPs in tested region of HR-related genes.

## Results

The performed analysis aimed at determination of significance of new genetic variants as breast cancer risk factors. The study was carried out in a group of 94 breast cancer patients and in 500 disease-free controls. Microarray analysis identifies statistically significant correlations among SNPs localised on chromosomes 13 and 17. A pool of 3 SNPs mostly correlated with breast cancer risk was determined (see Table 3). These single nucleotide polymorphisms included: rs59004709 (chromosome 13, *BRCA2*), rs4986852 and rs1799950 (last two SNPs on chromosome 17, *BRCA1*). See Table 3 for the areas with genes at which the above-mentioned SNPs were localised.

**Table 3.**
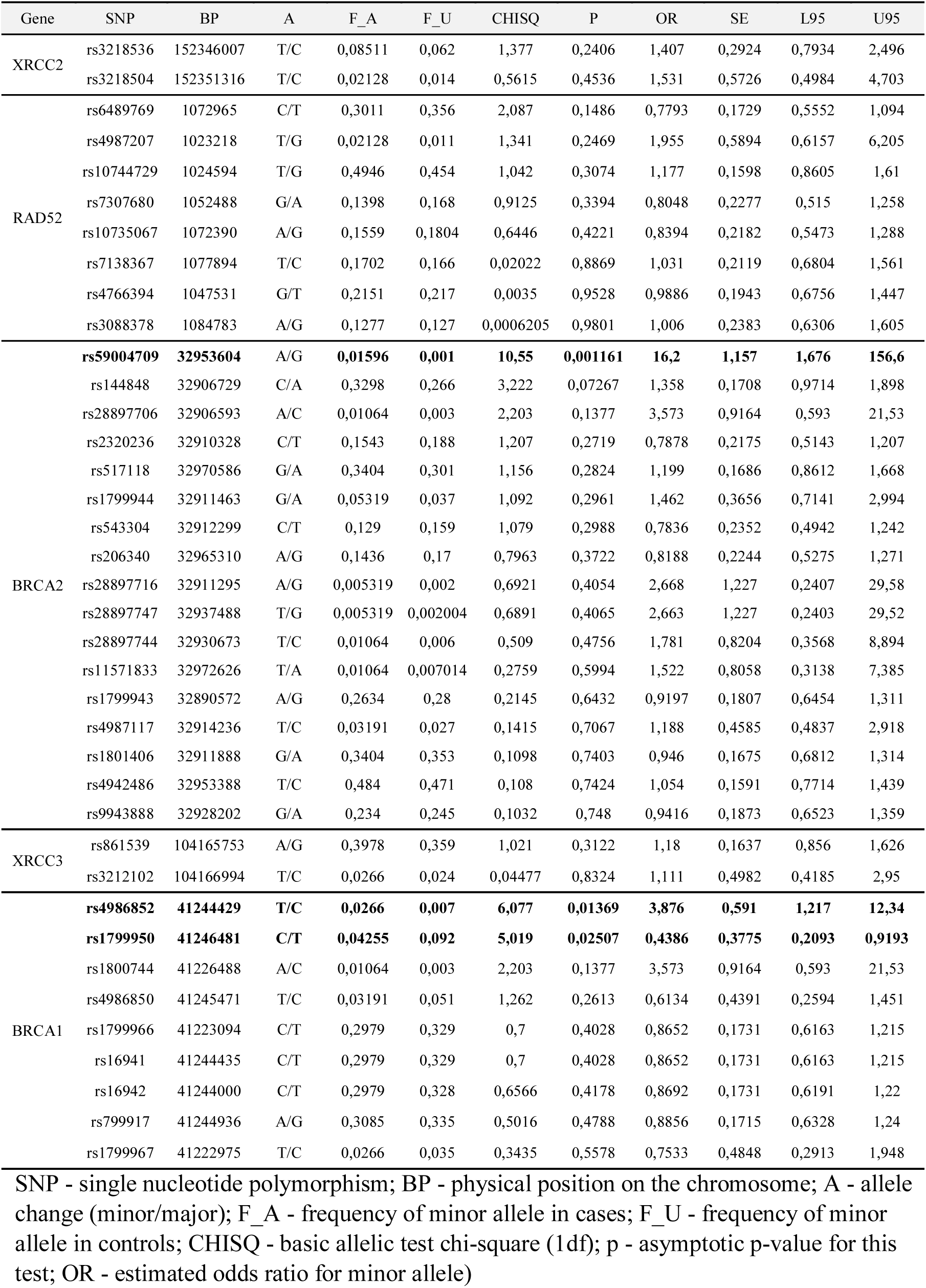
Case-control association analysis of HR-related gene SNPs.

| Gene | SNP | BP | A | F_A | F_U | CHISQ | P | OR | SE | L95 | U95 |
| --- | --- | --- | --- | --- | --- | --- | --- | --- | --- | --- | --- |
| XRCC2 | rs3218536 | 152346007 | T/C | 0,08511 | 0,062 | 1,377 | 0,2406 | 1,407 | 0,2924 | 0,7934 | 2,496 |
|  | rs3218504 | 152351316 | T/C | 0,02128 | 0,014 | 0,5615 | 0,4536 | 1,531 | 0,5726 | 0,4984 | 4,703 |
| RAD52 | rs6489769 | 1072965 | C/T | 0,3011 | 0,356 | 2,087 | 0,1486 | 0,7793 | 0,1729 | 0,5552 | 1,094 |
|  | rs4987207 | 1023218 | T/G | 0,02128 | 0,011 | 1,341 | 0,2469 | 1,955 | 0,5894 | 0,6157 | 6,205 |
|  | rs10744729 | 1024594 | T/G | 0,4946 | 0,454 | 1,042 | 0,3074 | 1,177 | 0,1598 | 0,8605 | 1,61 |
|  | rs7307680 | 1052488 | G/A | 0,1398 | 0,168 | 0,9125 | 0,3394 | 0,8048 | 0,2277 | 0,515 | 1,258 |
|  | rs10735067 | 1072390 | A/G | 0,1559 | 0,1804 | 0,6446 | 0,4221 | 0,8394 | 0,2182 | 0,5473 | 1,288 |
|  | rs7138367 | 1077894 | T/C | 0,1702 | 0,166 | 0,02022 | 0,8869 | 1,031 | 0,2119 | 0,6804 | 1,561 |
|  | rs4766394 | 1047531 | G/T | 0,2151 | 0,217 | 0,0035 | 0,9528 | 0,9886 | 0,1943 | 0,6756 | 1,447 |
|  | rs3088378 | 1084783 | A/G | 0,1277 | 0,127 | 0,0006205 | 0,9801 | 1,006 | 0,2383 | 0,6306 | 1,605 |
| BRCA2 | **rs59004709** | **32953604** | A/G | **0,01596** | **0,001** | **10,55** | **0,001161** | **16,2** | **1,157** | **1,676** | **156,6** |
|  | rs144848 | 32906729 | C/A | 0,3298 | 0,266 | 3,222 | 0,07267 | 1,358 | 0,1708 | 0,9714 | 1,898 |
|  | rs28897706 | 32906593 | A/C | 0,01064 | 0,003 | 2,203 | 0,1377 | 3,573 | 0,9164 | 0,593 | 21,53 |
|  | rs2320236 | 32910328 | C/T | 0,1543 | 0,188 | 1,207 | 0,2719 | 0,7878 | 0,2175 | 0,5143 | 1,207 |
|  | rs517118 | 32970586 | G/A | 0,3404 | 0,301 | 1,156 | 0,2824 | 1,199 | 0,1686 | 0,8612 | 1,668 |
|  | rs1799944 | 32911463 | G/A | 0,05319 | 0,037 | 1,092 | 0,2961 | 1,462 | 0,3656 | 0,7141 | 2,994 |
|  | rs543304 | 32912299 | C/T | 0,129 | 0,159 | 1,079 | 0,2988 | 0,7836 | 0,2352 | 0,4942 | 1,242 |
|  | rs206340 | 32965310 | A/G | 0,1436 | 0,17 | 0,7963 | 0,3722 | 0,8188 | 0,2244 | 0,5275 | 1,271 |
|  | rs28897716 | 32911295 | A/G | 0,005319 | 0,002 | 0,6921 | 0,4054 | 2,668 | 1,227 | 0,2407 | 29,58 |
|  | rs28897747 | 32937488 | T/G | 0,005319 | 0,002004 | 0,6891 | 0,4065 | 2,663 | 1,227 | 0,2403 | 29,52 |
|  | rs28897744 | 32930673 | T/C | 0,01064 | 0,006 | 0,509 | 0,4756 | 1,781 | 0,8204 | 0,3568 | 8,894 |
|  | rs11571833 | 32972626 | T/A | 0,01064 | 0,007014 | 0,2759 | 0,5994 | 1,522 | 0,8058 | 0,3138 | 7,385 |
|  | rs1799943 | 32890572 | A/G | 0,2634 | 0,28 | 0,2145 | 0,6432 | 0,9197 | 0,1807 | 0,6454 | 1,311 |
|  | rs4987117 | 32914236 | T/C | 0,03191 | 0,027 | 0,1415 | 0,7067 | 1,188 | 0,4585 | 0,4837 | 2,918 |
|  | rs1801406 | 32911888 | G/A | 0,3404 | 0,353 | 0,1098 | 0,7403 | 0,946 | 0,1675 | 0,6812 | 1,314 |
|  | rs4942486 | 32953388 | T/C | 0,484 | 0,471 | 0,108 | 0,7424 | 1,054 | 0,1591 | 0,7714 | 1,439 |
|  | rs9943888 | 32928202 | G/A | 0,234 | 0,245 | 0,1032 | 0,748 | 0,9416 | 0,1873 | 0,6523 | 1,359 |
| XRCC3 | rs861539 | 104165753 | A/G | 0,3978 | 0,359 | 1,021 | 0,3122 | 1,18 | 0,1637 | 0,856 | 1,626 |
|  | rs3212102 | 104166994 | T/C | 0,0266 | 0,024 | 0,04477 | 0,8324 | 1,111 | 0,4982 | 0,4185 | 2,95 |
| BRCA1 | **rs4986852** | **41244429** | **T/C** | **0,0266** | **0,007** | **6,077** | **0,01369** | **3,876** | **0,591** | **1,217** | **12,34** |
|  | **rs1799950** | **41246481** | **C/T** | **0,04255** | **0,092** | **5,019** | **0,02507** | **0,4386** | **0,3775** | **0,2093** | **0,9193** |
|  | rs1800744 | 41226488 | A/C | 0,01064 | 0,003 | 2,203 | 0,1377 | 3,573 | 0,9164 | 0,593 | 21,53 |
|  | rs4986850 | 41245471 | T/C | 0,03191 | 0,051 | 1,262 | 0,2613 | 0,6134 | 0,4391 | 0,2594 | 1,451 |
|  | rs1799966 | 41223094 | C/T | 0,2979 | 0,329 | 0,7 | 0,4028 | 0,8652 | 0,1731 | 0,6163 | 1,215 |
|  | rs16941 | 41244435 | C/T | 0,2979 | 0,329 | 0,7 | 0,4028 | 0,8652 | 0,1731 | 0,6163 | 1,215 |
|  | rs16942 | 41244000 | C/T | 0,2979 | 0,328 | 0,6566 | 0,4178 | 0,8692 | 0,1731 | 0,6191 | 1,22 |
|  | rs799917 | 41244936 | A/G | 0,3085 | 0,335 | 0,5016 | 0,4788 | 0,8856 | 0,1715 | 0,6328 | 1,24 |
|  | rs1799967 | 41222975 | T/C | 0,0266 | 0,035 | 0,3435 | 0,5578 | 0,7533 | 0,4848 | 0,2913 | 1,948 |

SNP - single nucleotide polymorphism; BP - physical position on the chromosome; A - allele change (minor/major); F\_A - frequency of minor allele in cases; F\_U - frequency of minor allele in controls; CHISQ - basic allelic test chi-square (1df); p - asymptotic p-value for this test; OR - estimated odds ratio for minor allele)

The potential relationship between *XRCC2*, *XRCC3, RAD51, RAD52, BRCA1* and *BRCA2* SNPs genotype distribution and clinical features of breast cancer patients was investigated. However, the current study failed to show any correlation of analysed SNPs with: tumour size, grade or lymph node status. Nor were DNA repair genes’ polymorphisms related to the patients’ estrogen (ER) and progesterone (PR) receptors status (p > 0.05).

## Discussion

Defective DNA repair system has been already suggested as a predisposing factor in familial and sporadic breast cancer [22, 23]. Furthermore, it has been demonstrated that SNPs of DNA repair genes may independently contribute to increased risk of breast carcinogenesis due to the disorders they provoke in maintaining genome integrity [24-27].

In this study the authors’ interest was focused on the analysis of SNPs of HR-related genes in a group of breast cancer patients and in healthy controls.

RAD51 homolog (RecA homolog, E. coli) (S. cerevisiae) plays an important role in HR through direct interaction with XRCC2 (X-ray repair cross-complementing group 2), XRCC3 (X-ray repair cross-complementing group 3), BRCA1/2 (breast cancer-1/2) and other DNA repair proteins. Common genetic variants in genes involved in DNA DSBs repair are plausible candidates for low-penetrance breast cancer susceptibility.

Such a genetic variation of *RAD51* has already been proposed to coincide – among others – with breast cancer [28-33]. Current literature already confirms the significance of *RAD51*-G135C polymorphism in breast cancer risk [32, 33]. Moreover, some reports determine *XRCC2* and *XRCC3* variation as related to increased risk of this malignancy [34, 35, 18, 35-38]. Also, it has been earlier shown that the combined Arg188His-*XRCC2*/Thr241Met-*XRCC3*/135G/C-*RAD51* genotype increased the risk of breast cancer in Polish population [33].

Recent studies on versatile populations have provided first epidemiological evidence supporting the association between *BRCA1/2* repair genes’ variants and breast cancer development [39-41]. The association between the triple-negative phenotype and breast cancer harboring germline mutations in the *BRCA1* gene has been previously well described. Mutations in this crucial gene confer an approximately 80% lifetime risk of breast cancer among carriers [42, 43]. In Polish women several studies confirm the significance of *BRCA1* and *BRCA2* genes polymorphism in the risk of breast carcinoma [44, 45]. However, the reported studies’ results have rather been inconsistent [46, 47].

The abovementioned results of SNPs analysis (as well as those discussed in the Introduction) reveal the existence of certain links with breast cancer. Learning their detailed structure may contribute to the development of new therapeutic strategies.

Genetic analysis presented in this report revealed statistically significant correlations between breast cancer and 3 not yet described SNPs. The polymorphisms were found within the areas of the *BRCA1/2* genes for which − at least basing on available literature data − no explanation could be found for their potential role in breast cancer pathogenesis. It should be emphasised that the study was carried out on rather limited groups and thus the obtained results should be approached as preliminary. We strongly believe that genetic analysis performed on much larger groups may result in a better understanding of the pathology of breast cancer. Taking into account how scarce is the up-to-date literature in analysing correlation between SNPs and breast cancer, one may conclude that the abovementioned results introduce an innovative quality into breast cancer research on an international scale. However, we must emphasise again that the quantitative insufficiency of our groups allows us to consider it to be only a pilot study revealing just some preliminary piece of information on genetic markers in breast cancer. In conclusion, authors encouraged by such promising results intend to continue this study.

## Acknowledgments

This study was supported by the Polish POIG grant 01.01.02-10-005/08 TESTOPLEK from the European Regional Development Fund.

## Authors’ contributions

Conceived and designed the experiments: DS, BS. Performed the experiments – case group: MSK, MS. Case group design and collect: HR, BS, MZ. Performed the experiments – control group: MSK, MS, EK, ASG. Analysed case-control data: MS, MSK, DS. Contributed reagents/materials/analysis tools DS. Contributed to the writing of manuscript: BS, HR, DS. All authors approved the final manuscript.

## Compliance with Ethical Standards

This study was supported by the Institute of Polish Mother’s Memorial Hospital, Lodz, Poland from the Statutory Development Fund. All procedures performed in studies involving human participants were in accordance with the ethical standards of the institutional and/or national research committee and with the 1964 Helsinki declaration and its later amendments or comparable ethical standards.

## Ethics approval

All the study participants gave a informed written consent. A formal consent was also issued by the Bioethical Committee of the Institute of the Polish Mother’s Memorial Hospital in Lodz (Approval number, 10/2012) and Review Board at the University of Lodz (Approval number 7/KBBN-UŁ/II/2014).

## Availability of data and material

The datasets supporting the conclusions of this article are included within the article.

